# Personality and behavioral syndromes in two species of fruit bats (Chiroptera: Phyllostomidae)

**DOI:** 10.1101/2023.11.23.568433

**Authors:** Pedro Henrique Miguel, Augusto F. Batisteli, Ariovaldo P. Cruz-Neto

## Abstract

Personality indicates consistency in individual behavioral responses across different contexts, and different personality traits may be correlated in behavioral syndromes. Studies on personality have important consequences for conservation ecology and, despite the ecological relevance of fruit bats, rare studies have tested the existence of personality and behavior syndromes in this group. In this context, this study aims to test whether the Neotropical fruit bats *Artibeus lituratus* and *Carollia perspicillata* present (1) individual repeatability (i.e. personality) related to the behaviors: activity, aggressiveness and boldness and (2) correlation between these personality axes, constituting behavioral syndromes. For each species, 27 adult males were captured and immediately placed in cloth bags individually, and we measured aggressiveness as the time they struggled in the bag within a 180-second interval. Bats were then kept in individual cages in a climate-controlled room for 48-h, after which we filmed them for 30 min in a flight tent as an activity test. In the boldness test, we quantified the latency to each animal to start feeding in front of an observer, assuming that bolder individuals had lower latency to feed. All tests were repeated after 48-h to analyze repeatability. We found high individual repeatability of aggressiveness, activity and boldness for both species, but different behavioral syndromes for each of them. For *C. perspicillata*, the three behaviors were correlated to each other, with the most aggressive individuals being bolder and more active. For *A. lituratus*, aggressiveness and boldness were also positively correlated, but activity did not correlate with the other two behaviors. Considering these different syndromes, our results suggest that *C. perspicillata* has less variation in behavioral profiles than *A. lituratus* probably as a result of species-specific selective pressures. The existence of behavioral syndromes in these frugivorous bats contributes to understanding the importance of individual variation on the ecological functions provided by seed dispersers.

---

Animal personality can be defined as the consistency over time (i.e. repeatability) of individual behavioral responses in different contexts (RÉALE et al., 2007; SIH et al., 2004). Different dimensions of animal behavior can be studied in isolation and still be referred to as a continuum of a given personality axis when showing repeatability at individual level (CAREAU; GARLAND, 2012; SIH; BELL, 2008). Because behavioral decisions affect fitness via process as resource acquisition and predator avoidance, animal personality traits are potentially under evolutive pressures through natural selection and have important ecological consequences (DINGEMANSE et al., 2004; SIH et al., 2004; ROBERTS; SHERRATT, 2002). For instance, bolder and more active individuals play a crucial role in dispersing and establishing new populations (DOHM, 2002; HAYES; JENKINS, 1997), increasing genetic diversity and promoting the long-term persistence of populations at regional scale (DINGEMANSE et al., 2002; MINDERMAN et al., 2009). Moreover, animal personality is linked to wildlife conservation as it affects the way species cope with environmental changes (CAREAU; GARLAND, 2012; DINGEMANSE et al., 2004; SIH et al., 2004), because certain personality traits can endow animals with greater flexibility and adaptability to thrive in the face of these ongoing environmental fluctuations (CAREAU; GARLAND, 2012; DINGEMANSE et al., 2004; SIH et al., 2004).

When two or more personality traits are correlated within a population, they constitute a behavioral syndrome (SIH et al., 2004; SIH; BELL, 2008). Although some specific personality traits or behavioral syndromes are better suited to certain contexts than others, fluctuations in environmental conditions should promote the coexistence of a range of personality types within populations (SIH et al., 2012; NICOLAUS et al., 2016). Understanding these individual differences is important to predict their potential impact on individual fitness and behavior selection (STAMPS; GROOTHUIS, 2010).

A classical axis of animal personality is the boldness/shyness gradient, which reflects the propensity of an individual face a risky situation, usually measured by their reaction to the proximity of predators or humans (CHAPMAN et al., 2010; STAMPS, 2007; WILSON et al., 1993). Boldness is therefore associated with the capacity to use less favorable habitat patches such as those with higher predation risk, because bolder individuals cope better with these challenges (COTE et al., 2010; DINGEMANSE et al., 2003; FRASER et al., 2001; STAMPS, 2007).

The capacity of moving between habitat patches is linked to another aspect of animal personality, the activity gradient, primarily associated with individual propensity for locomotion and movement in the environment (DINGEMANSE et al., 2002; REALE et al., 2007; VERBEEK et al., 1994). This concept is most prominently related to spatial movement in the wild, reflecting a tendency where individuals exhibiting higher activity levels in laboratory experiments often display similar tendencies in natural field settings (see COTE et al. 2010 for a review). Furthermore, compelling evidence indicates correlations between activity levels and various ecological factors, including individual dispersal distance (DINGEMANSE et al., 2003; FRASER et al., 2001; HOSET et al., 2010), seasonal dispersal patterns (CHAPMAN et al., 2011; VAN OVERVELD et al., 2014), and home range sizes (MINDERMAN et al., 2009; VAN OVERVELD et al., 2011). Hence, activity level represents an informative personality trait for investigating individual and population-level processes in the fields of behavioral and movement ecology (COTE et al., 2010; SIH et al., 2012).

A third personality axis, the docility/aggression gradient, relates to the individual’s reaction to capture and manipulation, a response mediated by glucocorticoid hormones (BIRO; STAMPS, 2008; CAREAU et al., 2015). It is commonly used as a measure of self-preservation, which has evolved due to the intricate trade-off between longevity and reproductive success (RÉALE et al., 2007; VAN OERS; SINN, 2013; ZWOLAK; SIH, 2020). Despite the relevance of each of these personality traits per se, the interplay between them and their impact on individual behavior can provide valuable insights into the complex dynamics of personality in animals. For instance, more aggressive individuals competing for scarce food resources, tend to perform better in intraspecific conflicts but may allocate more energy resources to offspring maintenance, potentially leading to shorter lifespans compared to more docile and vulnerable individuals (NICOLAUS et al., 2016).

The investigation of personality syndromes encompassing these three traits is of paramount importance for conservation (RÉALE et al., 2007; RÉALE et al., 2009; MACKINLAY; SHAW 2023). Investigating the potential syndromes involving docility/aggression, risk propensity, and activity level, holds the potential to yield profound insights into the mechanisms by which personality shapes different behavioral profiles within populations (CAREAU; GARLAND 2012; RÉALE et al., 2007; SIH et al., 2012). For instance, a syndrome where heightened aggression coincides with a greater willingness to take risks in resources acquisition leads to distinct behavioral patterns that impact an individual’s overall fitness (RÉALE et al., 2007; RÉALE et al., 2009; ZWOLAK; SIH, 2020). Syndromes like these have the potential to provide invaluable insights into the correlated manifestation of personality traits and their subsequent consequences (CAREAU; GARLAND 2012; DINGEMANSE et al., 2004; RÉALE et al., 2007). However, despite the importance of studying personality and syndromes, it’s worth noting that some groups, as is the case of bats, remain underrepresented in these studies.

Bats serve as valuable models for the study of animal personality and behavioral syndromes, as their individual behavioral traits can impact several ecological processes, including resource use, predation dynamics, pollination and seed dispersal, ultimately contributing to ecosystem maintenance (MUYLAERT et al. 2017; SIMMONS; CONWAY, 2003). Understanding the role of personality in frugivorous animals, such as bats, is particularly important, as it may have profound implications for seed dispersal, potentially resulting in consequences such as the decision-making process regarding resource consumption in specific locations (AZEVEDO; YOUNG, 2021; ZWOLAK; SIH, 2020). Personality traits and behavioral syndromes in bats potentially underline various decisions related to frugivory, which may lead to trade-offs related to the quality of seed dispersal by individuals with diverse behavioral tendencies (DINGEMANSE et al., 2004; SIH et al., 2012; ZWOLAK; SIH, 2020).

Bats, representing around 20% of all mammals on the planet, exhibit significant ecological and morphological diversity (SIMMONS; CONWAY, 2003). Among the approximately 288 species of Neotropical bats, approximately 83 are fruit consumers and seed dispersers (GORRESEN; WILLIG, 2004; MICKLEBURGH et al., 2002; SIMMONS, 2005), 28 of them found in Brazil (FABIAN et al., 2008). *Artibeus lituratus* and *Carollia perspicillata* (Phyllostomidae) are important seed dispersers abundant in preserved areas, small forest patches, and disturbed rural and urban environments (MUYLAERT et al., 2017; BALLESTEROS; RACERO-CASARRUBIA 2012, JARA-SERVÍN et al. 2017, NUNES et al. 2017). However, it’s worth noting that the few personality studies in bats are most focused on insectivorous species from temperate regions as models (MENZIES et al., 2013; WEBBER; WILLIS, 2020; BOYER et al., 2020), while studies on frugivorous bats are rare (e.g. HARTEN et al., 2021). Inhabiting the mentioned environmental disturbance gradient, *A. lituratus* and *C. perspicillata* are expected to exhibit a marked variation in personality traits that enhances that behavioral flexibility at population level. Exploring the personality of frugivorous animals, especially those capable of thriving in degraded environments, can yield valuable insights for bat conservation and forest restoration efforts.

In this context, this study aimed to test whether the frugivore bats *A. lituratus* and *C. perspicillata* present (1) individual repeatability related to the behaviors: activity, aggressiveness/docility and boldness/shyness, indicating they are personality traits (2) correlation between these personality axes, constituting behavioral syndromes. Our first hypothesis is that aggression, activity and boldness constitute personality traits for *A. lituratus* and *C. perspicillata* as in other bats (MENZIES et al., 2013; WEBBER; WILLIS, 2020). Our second hypothesis is that there are behavioral syndromes for these two species, composed by the correlation between boldness and activity because individuals that are bolder tend to exhibit more active behaviors (BIANCONI, 2004; SEKIAMA, 2001). Similarly, we predict a positive correlation between activity and aggressiveness as highly active individuals may engage more in aggressive behaviors as they encounter more frequently threats and competitors for resources and potential mates (BIRO; STAMPS, 2008; NICOLAUS et al., 2016; WOLF; WEISSING, 2012). We also expect to find a positive relationship between boldness and aggressiveness in the study species, where bolder bats would be more likely to exhibit aggressive behaviors (RÉALE et al., 2007; STAMPS, 2007; VAN OERS; SINN, 2013).

## MATERIAL AND METHODS

Bats were captured using mist nets from December 2021 to March 2022 at the Edmundo Navarro de Andrade State Forest, Rio Claro, SP (22°24’56 ”S; 47°31’17 ”W) and at the Federal University of São Carlos, São Carlos, SP (21°59’05 ”S; 47°52’50 ”W), Southeastern Brazil.

We captured 27 adult males of each species. After captured, each individual was placed in a cloth bag (25 cm high × 20 cm long) for the first measurement of aggressiveness (see below). All animals captured on each collection night were kept individually in cloth bags until the end of the collection period (23:00 h), when they were transported to the Laboratory of Animal Physiology at the Institute of Biosciences of the Universidade Estadual Paulista “Júlio de Mesquita Filho” at Rio Claro. In the lab, they were housed in individual cages (40 × 40 × 40 cm) and kept in a climatic chamber at 26 °C and photoperiod 12/12 h for 2 days before the other behavioral tests. Each test was conducted twice with each individual in an interval of 48 h between replicates to allow estimating individual repeatability of behaviors.

### Aggressiveness test

The handling bag test was performed as a measure of aggressiveness (CAREAU et al., 2015; MARTIN; REALE, 2008). This test consists of quantifying movement time, i.e., the time the animal struggles from the moment the animal is placed in the bag within a period of 180 seconds. Individuals who moved for a longer time were considered more aggressive (CAREAU et al., 2015; MARTIN; REALE, 2008). This test was performed right after the capture in the field and repeated in the lab after 48 h, before the beginning of the other experiments.

### Activity test

To measure activity level of the individuals, we adapted the open field test (MARTIN; RÉALE, 2008; MINDERMAN et al., 2009). For this, we built an arena measuring 230 × 110 × 110 cm (length × height × width), with the sides, top and bottom covered with PVC screen (mesh 5 × 7 mm) and transparent plastic on the front side, so that the animals’ activities could be recorded using a Sony Handycam HDR-CX405 camcorder positioned 2.5 m from this side. This arena was built in a closed room free of weathering and external interference, with temperature varying between 22°C to 26°C.

We individually placed each bat in this arena after 6:00 pm, when these animals begin their activities in the wild, and filmed them for 30 minutes. We assume animals that moved longer as more active (BREHM et al., 2019; METTKE-HOFMAN et al., 2006). After this test, the animals returned to the individual cage, where the boldness test described below was performed.

### Boldness test

The boldness test was performed right after the activity test in another closed room, free from weathering and external interference. For this test, the bats were conditioned in individual cages (40 × 40 × 40 cm) and fed in a round plastic bowl (5 cm in diameter) with papaya (*Carica papaya*). After introduce the bowl in the cage, the same researcher (PHM) stand up 30 cm distant from the cage, which was positioned 1.5 m above ground, and recorded the latency for the animal start eating. We assumed that individuals who took less time to start eating were bolder. After the first and second replicate of the boldness test, bats were weighed to the nearest 1 g and then returned to the climate-controlled room.

### Statistical analysis

In order to assess the existence of personalities, we performed repeatability tests using the ’rptGaussian’ function in the ’rptR’ package (BREHM et al., 2019; STOFFEL et al., 2017). This function uses linear mixed effects modeling to provide a repeatability estimate that ranges from 0 to 1 and its confidence interval is calculated using parametric bootstrapping and Bayesian method. The method tests whether the behavioral responses are consistent at the individual level, that is, whether the repeatability estimate differs from zero, through a likelihood ratio test (STOFFEL et al., 2017). In all models, we set body mass, site (Rio Claro and São Carlos) and trial (first vs. second test) as independent variables and defined 1000 iterations for parametric bootstrapping. To assess the effect of same variables in behavioral responses, we rebuild the same models using the packeges ’lme4’ (BATES et al., 2007) and ’lmerTest’ (KUZNETSOVA et al., 2015).

To analyze whether the personalities axes formed syndromes (correlation between personality axes), we used Pearson’s correlation. We used the average of the two replicates as a single measurement for each individual and behavioral test and tested the correlation of pairwise combinations of the three behaviors studied. To enhance the comprehension of the biological meaning of the boldness measurement, latency to feed was multiplied by −1 (e.g. zero latency is the highest boldness value) when addressing its relationship with aggressiveness and activity in correlation tests.

All analyzes were conducted in the R software (R CORE TEAM, 2022). Repeatability estimates, denoted as R, are followed by their confidence intervals as CI [minimum; maximum], and other values are presented as mean ± standard deviation unless otherwise noted.

### Ethical Note

This study received ethical approval from the Coordination of the Animal Use Ethics Committee (CEUA) at the Bioscience Institute of UNESP (Universidade Estadual Paulista) in Rio Claro, São Paulo, Brazil (number 18/2021) and adhered to the ASAB/ABS guidelines. Following the completion of the study, all captured bats were released at their original capture location

## RESULTS

Aggressiveness was not affected by body mass, study sites or trials for both species (Table 1, Table 2). We found repeatability of aggressiveness for *C. perspicillata* (R = 0.651; CI = 0.395–0.837; p < 0.00; Fig. 1A) and for *A. lituratus*, (R = 0.704; CI = 0.478–0.866; p < 0.001; Fig. 1B).

**Fig. 1.**
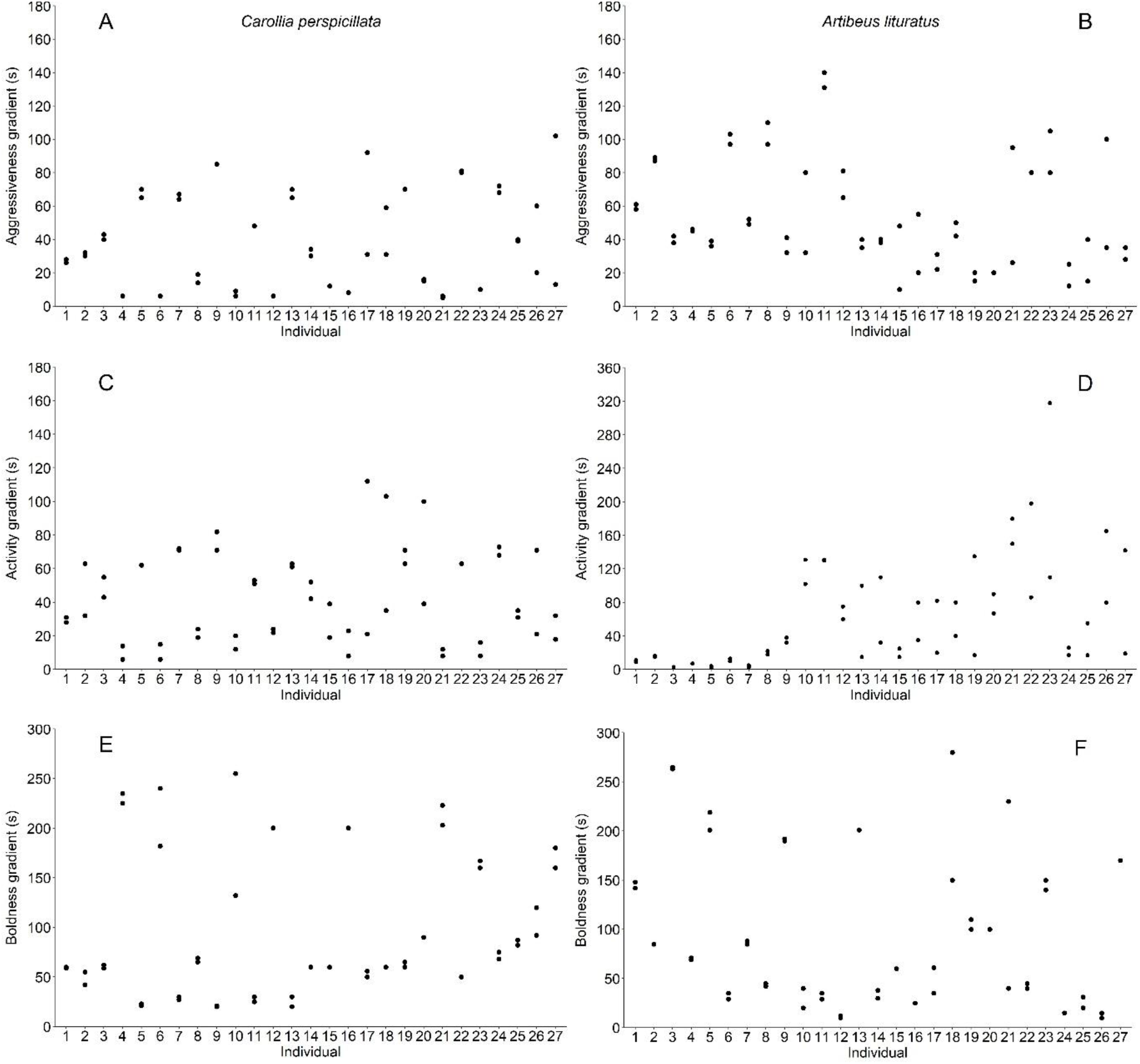
Individual variation in aggressiveness, activity and boldness in *Carollia perspicillata* (left, A, C and E) and *Artibeus lituratus* (right, B, D, F) measured respectively by the time struggling in the bag-handling test, time moving in the flight tent and latency to start feeding in face of an observer, when latency 0 corresponds to the highest boldness value.

**Table 1.**
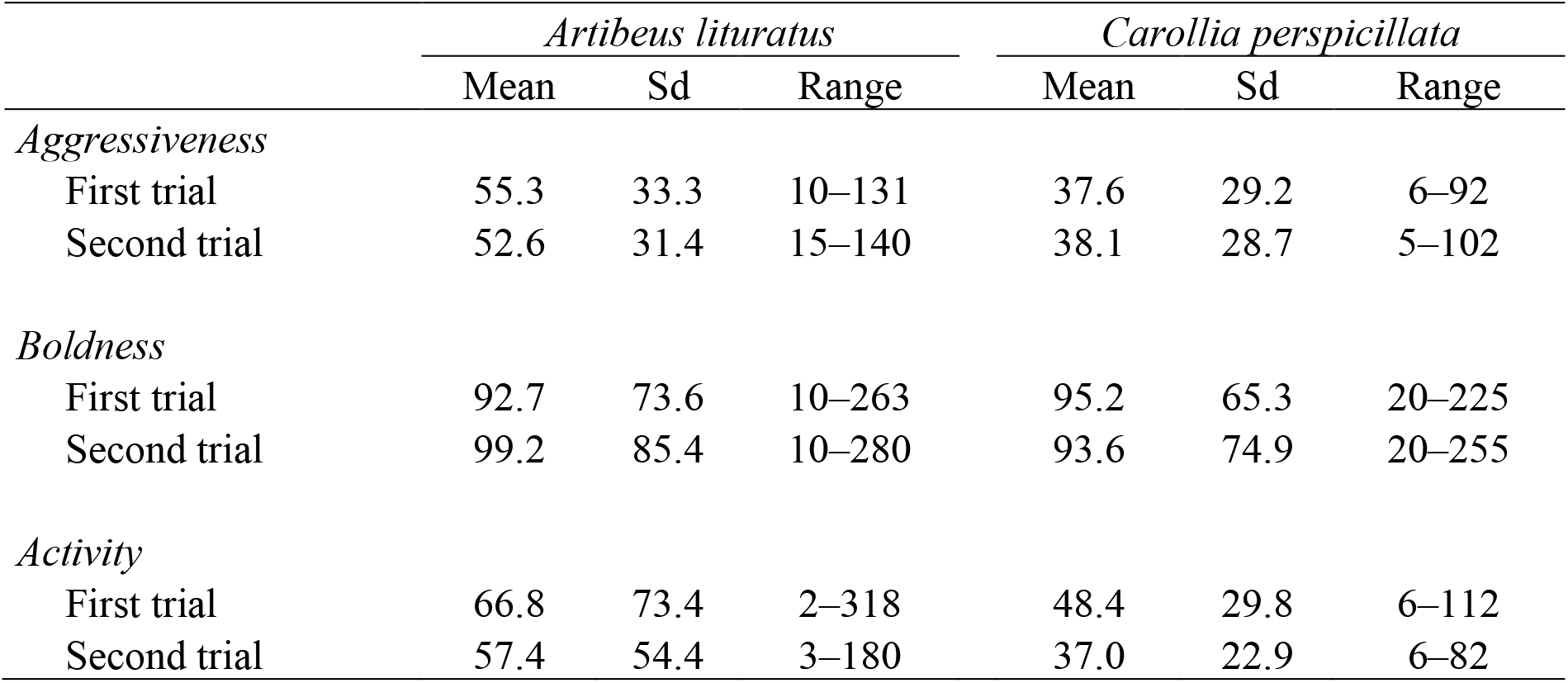
Variation of aggressiveness, boldness, and activity levels in *Artibeus lituratus* and *Carollia perspicillata* between two replicates of the tests, all measured in seconds as time struggling during handling-bag tests, latency to feed near an observer and time moving inside a flight tent, respectively.

|  | *Artibeus lituratus* |  |  | *Carollia perspicillata* |  |  |
| --- | --- | --- | --- | --- | --- | --- |
|  | Mean | Sd | Range | Mean | Sd | Range |
| *Aggressiveness* |  |  |  |  |  |  |
| First trial | 55.3 | 33.3 | 10–131 | 37.6 | 29.2 | 6–92 |
| Second trial | 52.6 | 31.4 | 15–140 | 38.1 | 28.7 | 5–102 |
| *Boldness* |  |  |  |  |  |  |
| First trial | 92.7 | 73.6 | 10–263 | 95.2 | 65.3 | 20–225 |
| Second trial | 99.2 | 85.4 | 10–280 | 93.6 | 74.9 | 20–255 |
| *Activity* |  |  |  |  |  |  |
| First trial | 66.8 | 73.4 | 2–318 | 48.4 | 29.8 | 6–112 |
| Second trial | 57.4 | 54.4 | 3–180 | 37.0 | 22.9 | 6–82 |

**Table 2.**
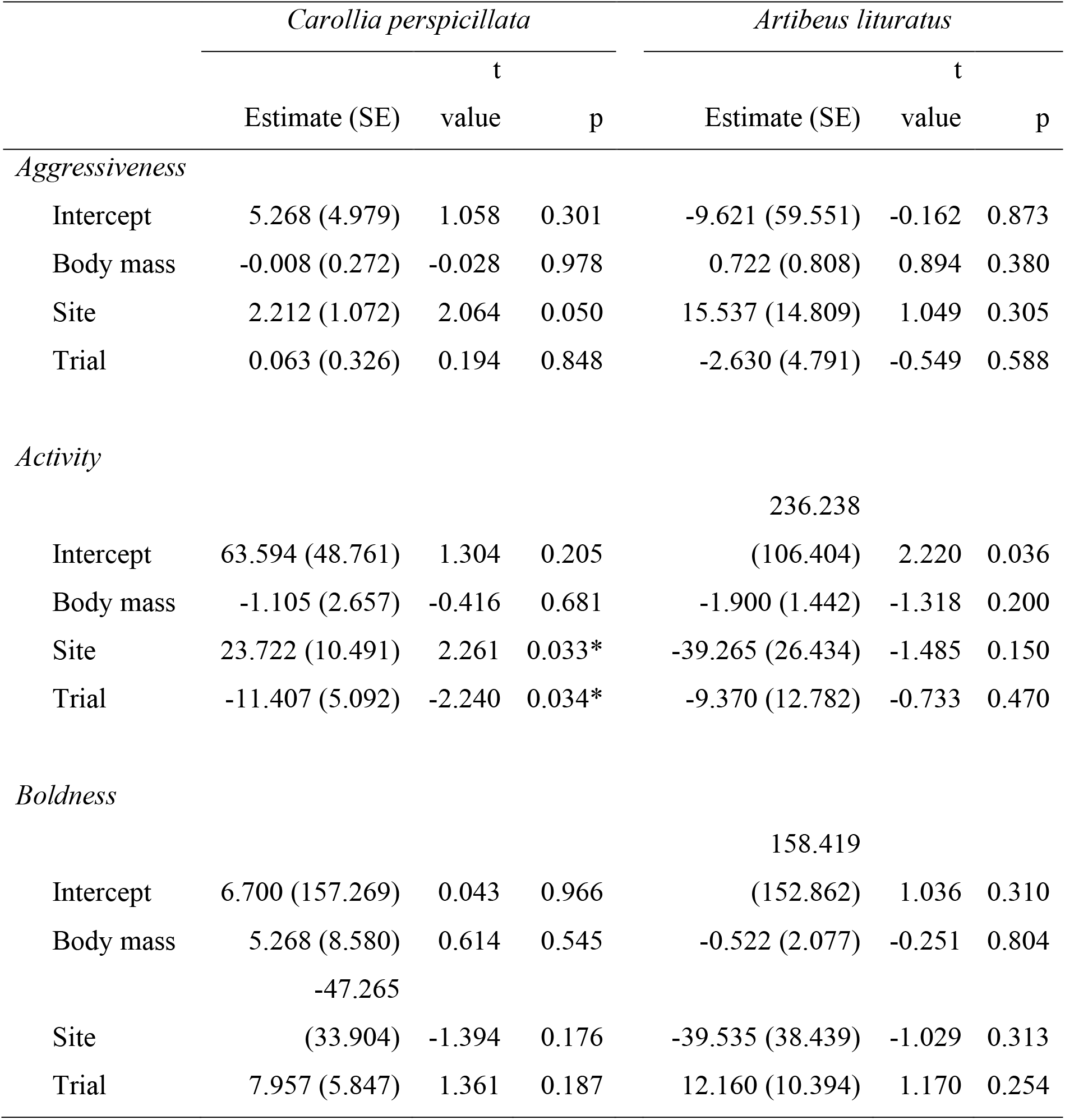
Results of linear mixed effects models assessing the effect of body mass, site (Rio Claro and São Carlos) and trial (first and second test) on individual responses of two Neotropical frugivorous bats (*Carollia perspicillata* and *Artibeus lituratus*) during behavioral tests of aggressiveness, boldness and activity. SE: Standard error; *: significant p-values at α = 0.05.

Activity level differed between sites and trials only for *C. perspicillata*, being lower for individuals captured in São Carlos and in the second trial of the test (Table 1, Table 2). For both species, body mass did not affect activity level. Activity level was repeatable for *C. perspicillata* (R = 0.461; CI = 0.123–0.749; p = 0.009; Fig. 1C) and for *A. lituratus* (R = 0.441; CI = 0.123–0.717; p = 0.013; Fig. 1D).

Finally, boldness was not affected by body mass, study sites or trials for both species (Table 1, Table 2). Boldness was repeatable for *C. perspicillata*, (R = 0.919; CI = 0.846–0.969; p < 0.001; Fig 1E) and for *A. lituratus* (R = 0.811; CI = 0.620–0.926; p < 0.001; Fig.1F).

There was a positive correlation between boldness and aggressiveness for *C. perspicillata* (r = 0.770; p < 0.001; Fig. 2A) and for *A. lituratus* (r = 0.419; p = 0.020; Fig. 2B). Activity level was also positively correlated with boldness for *C. perspicillata* (r = 0.802; p < 0.001; Fig. 2C), but not for *A. lituratus* (r = 0.158; p = 0.430). Similarly, activity was correlated with aggressiveness for *C. perspicillata* (r = 0.876; p < 0.001; Fig. 2D), but not for *A. lituratus* (r = 0.365; p = 0.060).

**Fig. 2.**
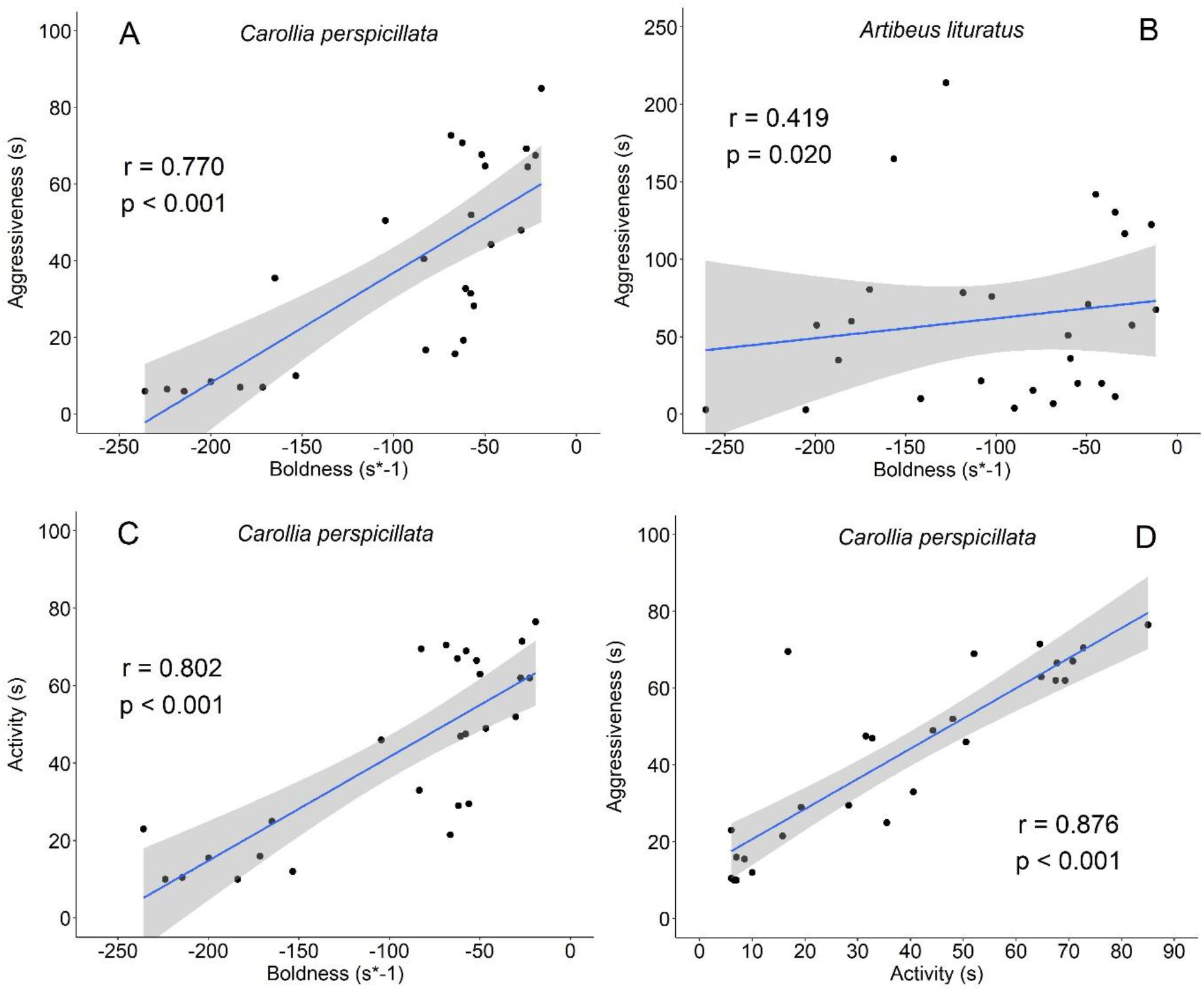
Correlations between boldness and aggressiveness behaviors for *Carollia perspicillata* (A) and *Artibeus lituratus* (B), activity and boldness (C) and activity and aggressiveness (D) for *Carollia perspicillata*. To enhance the comprehension of the biological meaning of boldness, latency to feed was multiplied by −1 (e.g. zero latency is the highest boldness value). The straight line indicates the trendline, and the hatched area indicates its 95% standard error.

## DISCUSSION

The three behavioral axes studied showed repeatability at individual level for *A. lituratus* and *C. perspicillata*, corroborating our first hypothesis, so that we can consider them personality traits in both species. This result is in line previous studies on insectivorous bats (WEBBER; WILLIS, 2020; BOYER et al., 2020) and numerous evidences from other taxa (BELL et al., 2009), and reinforces that intraspecific variation in behavioral decisions has an expressive relevance to understand ecological and evolutive aspects of populations (MACKINLAY; SHAW 2023).

In general, the individual levels of the behaviors studied were not related to body mass and did not differ between sites and trials, indicating that intrinsic individual variation overlap these other aspects. The exception was the activity level of *C. perspicillata* individuals, which was slightly lower for individuals from São Carlos and in the second than in the first trial. The different baseline levels of activity for the two study sites probably arise from local factors beyond the scope of this study, but it draw attention to the importance of considering the spatial structuring of individual behavioral responses even for widely used personality traits (VAN DONGEN et al., 2010; MACKINLAY; SHAW 2023). The slightly lower activity level during the second test in *C. perspicillata* might indicate some consequence of captivity or habituation to testing conditions, but it should have occurred in *A. lituratus* as well, which was not the case. One could argue that *C. perspicillata* has a smaller body size than *A. lituratus* (18.5 ± 1.7 g vs 72.3 ± 7.3 g, respectively) and might have suffered a faster depletion of body condition during the captive period, but the body mass of the individuals remained the same between the first and the second trials. Whatever is the reason behind the lower activity of *C. perspicillata* individuals during the second trial, it did not compromise the consistency of this behavior at individual level, as denoted by its significant repeatability.

We found behavioral syndromes for both species studied, whereas previous studies on bats report mixed evidence supporting behavioral syndromes (WEBBER; WILLIS 2020; KUO et al., 2023) or not (BOYER et al., 2020; WANG et al., 2020). We highlight, however, that our second hypothesis was not fully supported as we found different behavioral syndromes for each species, given the correlation between boldness and aggressiveness for both, and the correlation of aggressiveness with activity and boldness only for *C. perspicillata*. The part of the behavioral syndrome common to the two model species was composed by boldness and aggressiveness, where the more aggressive the individuals, the bolder they were. One possible explanation for this involves responses to predators, where more bolder animals need to be more aggressive to thrive facing predators or competitors, while less daring (shy) animals tend to be evasive and avoid risks (DINGEMANSE; WOLF, 2010; LE COEURS et al., 2015; RÉALE et al., 2010).

*Carollia perspicillata* and *A. lituratus* exhibited a common correlation between boldness and aggressiveness, which can be attributed to the underlying mechanisms driving personality traits in animals (DOUGHERTY; GUILLETTE, 2018; WOLF; WEISSING, 2012). Boldness and aggressiveness are often linked as individuals with bolder personalities tend to risk more and display higher levels of aggression when face threats or competition (ZWOLAK; SIH, 2020; WOLF; WEISSING, 2012). This correlations is potentially advantageous in terms of resource acquisition, territory defense, and predator avoidance (MACKINLAY; SHAW, 2022; MOIRON et al., 2020; WOLF; WEISSING, 2012).

There were differences in the behavioral syndromes between the two species as activity was correlated to aggressiveness and boldness only in *C. perspicillata*. Potential factors related to this difference include some ecological and behavioral particularities of each species. *Carollia perspicillata*, being dominant in fragmented areas, may face higher predation risk due to the increased exposure in such habitats (ANDRADE et al., 2013; FLEMING; HEITHAUS 1981; LAURINDO et al., 2017). Consequently, higher activity levels would demand a selection for greater aggressiveness to enable these individuals to deal with potential threats associated with habitat fragmentation. In contrast, *A. lituratus* is dominant either in continuous forest or fragmented areas (ANDRADE et al., 2013; GALETTI; MORELLATO, 1994; MEDINA et al., 2007). The species demonstrated individual variation in activity levels independently of aggressiveness and boldness, which results in a greater flexibility of individual behavioral profiles. This high plasticity at population level may translate into greater adaptability, allowing *A. lituratus* to occupy habitat patches within a wide disruption degree, making it more resilient to environmental changes (ANDRADE et al., 2013; GALETTI; MORELLATO, 1994; MEDINA et al., 2007).

The observed differences in behavioral syndromes between the two species may also be associated to divergences in their foraging habits. *Artibeus lituratus* exhibits lower fidelity to specific locations and a broader foraging rang, influenced by food availability, particularly of *Ficus ssp.* (GALETTI; MORELLATO, 1994; MEDINA et al., 2007). In contrast, *C. perspicillata*, due to a lower flight capacity and specialized feeding preferences for *Piper* ssp., displays greater fidelity to specific foraging areas, favored by the edge effect in fragmented habitats (ANDRADE et al., 2013; FLEMING; HEITHAUS 1981; LAURINDO et al., 2017).

Our study revealed that two species of frugivorous bats present different behavioral syndromes, which shed light on the understanding of the ecological functions provided by each of them, since their behavioral profiles potentially modulate several stages of zoochoric seed dispersal, such as selecting the feeding location, dispersal distance and whether an animal returns to fetch the seed (BREHM et al., 2019; HUNTER JR et al., 2022; ZWOLAK; SIH, 2020). Moreover, by indicating that activity is repeatable in frugivorous bats, our results indicate a way by which a personality trait may affect seed dispersal quality at individual level, a neglected aspect of plant-animal interaction, which deserves further studies (ZWOLAK; SIH, 2020).

## CONCLUDING REMARKS

Our data indicate that behaviors related to aggressiveness, boldness and activity are linked to personality in *C. perspicillata* and *A. lituratus*, presenting relatively high degrees of repeatability at the individual level. We also found that some of these behaviors may be associated with each other forming species-specific behavioral syndromes. The structure of these behavioral syndromes suggests that *C. perspicillata* has less intraspecific variation in behavioral profiles than *A. lituratus*, since the three behavioral aspects studied are being selected apparently in an associated way in the former but not in the last. The behavior responsible for this difference in the composition of behavioral syndromes between species is activity, which may be linked to selective forces of divergent natures acting on each of them regarding movement ecology. The results of this study are particularly important to demonstrate the need to evaluate the biological performance of these species in different ecological scenarios from the perspective of personality and behavioral syndromes.

## Data Availability

Data that support the findings of this study are available from the corresponding author upon reasonable request.

## ACKNOWLEDGEMENTS

We would like to thank all members of the UNESP Conservation Physiology Laboratory (Lafa) for their help with field collections. PHM was supported by the Coordenação de Aperfeiçoamento de Pessoal de Nível Superior - Brasil (CAPES). AFB receives a postdoctoral grant from São Paulo Research Foundation (FAPESP #2020/12211-0). APCN was supported by a research grant from São Paulo Research Foundation (FAPESP # 2014/16320-7).

